# Transcriptome brings variations of gene expression, alternative splicing, and structural variations into gene-scale trait dissection in soybean

**DOI:** 10.1101/2023.07.03.545230

**Authors:** Delin Li, Qi Wang, Yu Tian, Xiangguang Lyv, Hao Zhang, Yinglu Sun, Huilong Hong, Huawei Gao, Yan-Fei Li, Chaosen Zhao, Jiajun Wang, Ruizhen Wang, Jinliang Yang, Bin Liu, Patrick S. Schnable, James C. Schnable, Ying-Hui Li, Li-Juan Qiu

## Abstract

Genome-wide association study (GWAS) identifies trait-associated loci, but due in part to slow decay of linkage disequilibrium (LD), identifying the causal genes can be a bottleneck. Transcriptome-wide association study (TWAS) addresses this by identifying gene expression-phenotype associations or integrating gene expression quantitative trait loci (eQTLs) with GWAS results. Here, we used self-pollinated soybean as a model to evaluate the application of TWAS in the genetic dissection of traits in plant species that exhibit slow LD decay. The first RNA-Seq analysis of a soybean diversity panel was conducted, which identified the genetic regulation of 29,286 genes. Different TWAS solutions were less affected by LD and robust with source of expression that identified known genes related to traits from different development stages and tissues. A novel gene named pod color *L2* was identified via TWAS and functionally validated by genome editing. Our introduction of the new exon proportion feature significantly improves the capture of expression variations resulting from structural variations and alternative splicing. As a result, the genes identified by our TWAS approach exhibited a diverse range of causal variations, including SNP, insertion/deletion, gene fusion, copy number variation, and alternative splicing. Using our TWAS approach, we identified genes associated with flowering time, including both previously known candidates and novel genes that had not been linked to this trait before, providing complementary insights with GWAS. In summary, this study supports the application of TWAS for candidate gene identification in species with low rates of LD decay.

## Introduction

Linking specific genes to the regulation of different plant traits is one of the fundamental goals of plant genetics. Successfully making these links provides insight into the underlying mechanisms responsible for plant growth, development, and adaptation. Genome-wide association study (GWAS) has been successful in identifying many such links based on correlations between genetic markers and phenotypic variation. However, identifying the specific causal genes responsible for marker-trait associations in GWAS can be challenging. One of the challenges of identifying the underlying causal genes responsible for marker trait associations is a particular problem in populations or species where linkage disequilibrium (LD) decays slowly across the genome (Flint-Garcia et al., 2003).

Measurements of transcript abundance across populations, like measurements of genetic markers, can identify links between genes and phenotypic variation. A number of approaches can be employed to identify genes responsible for variation in plant traits from transcriptome data, measurements of the abundance of the complete collection of RNA molecules expressed in a cell or tissue. Colocalization, which integrates both genetic marker and transcriptome data assesses the probability that the causal variants for gene expression quantitative trait loci (eQTLs) and GWAS-associated loci are identical on a locus-by-locus basis, using tools such as coloc (Giambartolomei et al., 2014) and eCAVIAR (Hormozdiari et al., 2016). eQTLs studies, by scanning association between variations of genotype and gene expression, have been conducted in most major crops or model plants (Jordan et al., 2007; Schadt et al., 2003; Wang et al., 2010; West et al., 2007). Transcriptome wide association study (TWAS) seek to establish association between transcript abundance – whether directly measured or inferred from *Cis*-regulatory variants -- and phenotypic variation directly. Profiling transcripts from equivalent tissues and environmental conditions for hundreds or thousands of samples remains both costly and logistically challenging. As a result, several methods for TWAS, including PrediXcan (Gamazon et al., 2015) and FUSION (Gusev et al., 2016), have been developed, which employed an expression reference panel to identify eQTLs and use these eQTLs to predict gene expression across a larger panel for which genetic marker data is available but directly measured transcript abundance is not. Another method is based on Mendelian randomization, which integrates eQTL and GWAS data from independent studies to identify candidate genes by using genetic variants as instrumental variables to infer causality between a risk factor, i.e., gene expression, and a trait outcome. SMR (Zhu et al., 2016) is one of the tools that can be used for Mendelian randomization analysis. The above approaches are able to reuse expression reference panels and eQTL analyses, once generated and conducted, to identify links between genes and traits in new populations (Anacleto et al., 2019; Chen et al., 2023; Li et al., 2020b; Yang et al., 2022). However, given sufficient effort and resources, it is also possible to conduct direct TWAS by measuring expression data across an entire association population that has also been phenotyped for a trait or traits of interest. As direct TWAS does not reply on genotype to infer expression levels it is less sensitive to LD decay – the associated issue of “multiple hits per locus” (Wainberg et al., 2019). The direct TWAS approach has been shown to identify genes missed by GWAS alone in plant species, including *Arabidopsis thaliana* (Li et al., 2021), *Brassica napus* (Tang et al., 2021), maize (*Zea mays*) (Kremling et al., 2019; Li et al., 2021; Lin et al., 2017; Zheng et al., 2020), and Sorghum (*Sorghum bicolor*) (Ferguson et al., 2021). However, direct TWAS studies to date have primarily focused on plants with either small genomes and/or rapid LD decay where linking associations to causal genes is less challenging.

Here we evaluate both inferred TWAS and direct TWAS approaches in a large diversity panel of soybean (*Glycine max* [L.] Merr.), a globally important oil seed crop where the characterization of natural genetic variation has been hindered by extremely slow linkage disequilibrium resulting from both population bottlenecks associated with domestication and improvement as well as a primarily self-pollinating reproductive strategy. Using transcript data measured via RNA-Seq from seedling shoots of a diverse soybean panel including wild (*Glycine soja* [Sieb. and Zucc.]), landrace, and improved soybean cultivars, we demonstrate that quantifying both structural variation (Scott et al., 2021) and alternative splicing variation (Gan et al., 2011; Wang et al., 2008) via “exon proportion” as part of a TWAS captures functional genetic variants which would be missed when simply considering overall transcript abundance. Our study demonstrates the robustness of TWAS using the expression data from seedling shoot, successfully identifying known genes associated with numerous qualitative and quantitative traits observed in different tissues and/or different developmental stages. TWAS in this population identified a putative causal gene for the soybean *L2* locus, which was subsequently validated via genome editing. TWAS identified both novel and known genes regulating variation in flowering time in soybean complementing GWAS results. Our findings suggest that the creation of large reference transcriptome panels will accelerate the process of linking genes to phenotypes in both soybean and other species with slow LD decay.

## Results

### RNA-Seq of wild, landrace, and improved soybeans

To obtain a diverse soybean transcriptome, we utilized a worldwide panel consisting of 70 wild, 304 landrace, and 248 improved soybean accessions for RNA-Seq (**Table S1**). Of these 622 accessions, 64 wild and 496 cultivars were previously genotyped via genome re-sequencing (Li et al., 2023b) (**Figure S1**). Seedling shoots in the V2 stage (**Figure 1A**) were collected from each soybean line, generating 5.1 Tb of raw RNA-Seq data. Each sample produced an average of 27.1 million pair-end reads of 2 x 150 bp, ranging from 6 Gb to 20 Gb (**Table S1**). After trimming, an average of 23.9 million pair-end reads were aligned to the annotated genes of the Wm82 V2 reference genome to measure gene expression levels and identify SNPs. 208,538 high-quality SNPs were retained and merged with previous re-sequencing SNPs (Li et al., 2023b), and 39,809 were novel SNPs from RNA-Seq.

**Figure 1.**
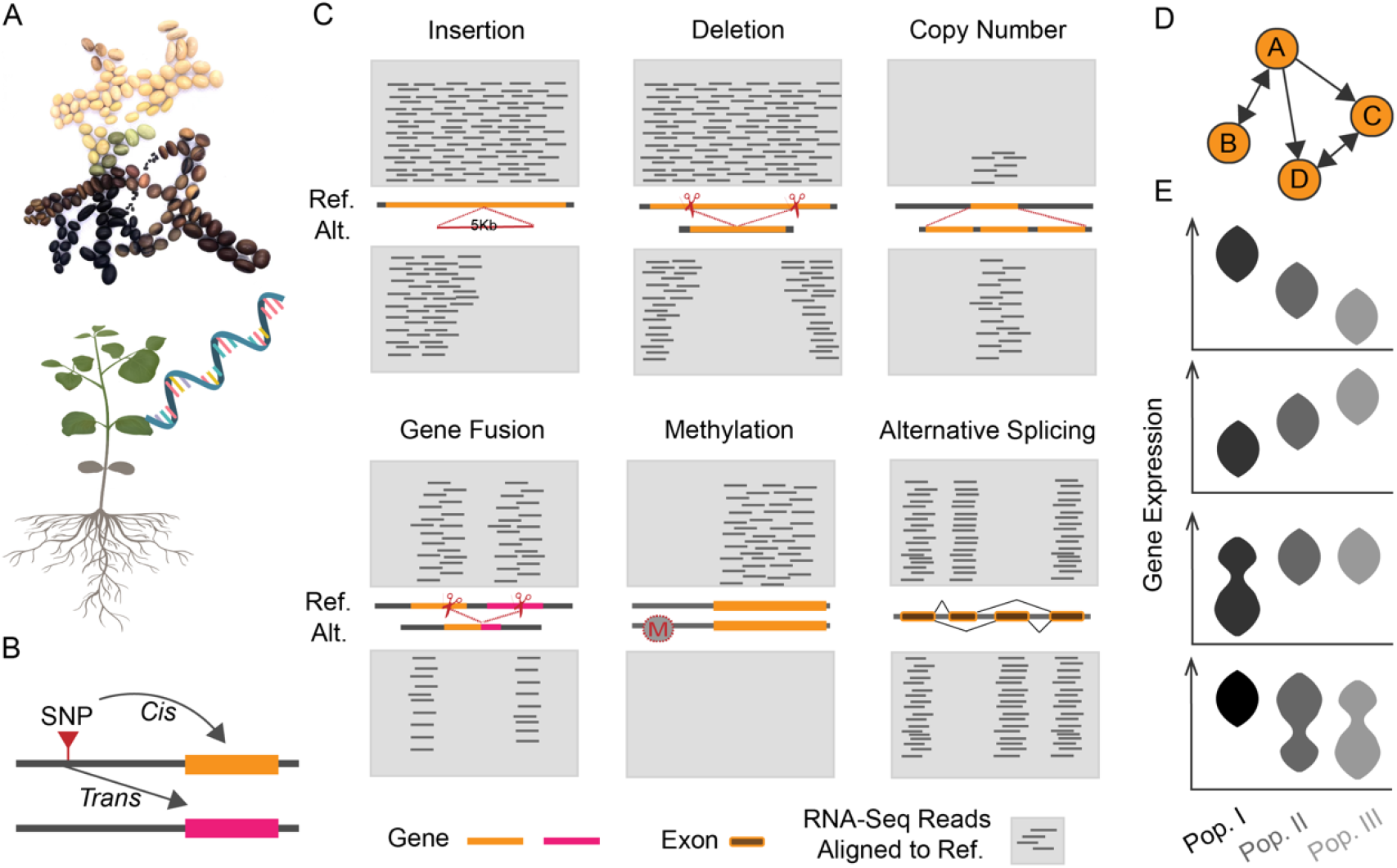
Information offered by the transcriptome. (**A**) In this study, transcriptomic data for 622 soybean accessions (70 wild ancestors, 304 landraces, and 248 improved lines) were generated using total RNA. The diversity of the panel is visualized with seed size and color in the archaic Chinese character for soybean, 菽. The green seedling shoot in the V2 stage was sampled for RNA-Seq, while the remaining part is depicted in grey. (**B**) The gene expression quantitative trait loci (eQTLs) could be identified by association studies between genetic variants (like SNPs) and gene expression. The eQTLs were divided into two categories, *Cis* and *Trans*, based on the distance between the variant and target gene, with *Cis* within 100Kb and *Trans* farther away. (**C**) Schematic diagrams of six types of variants that can be inferred from gene expression data but are typically not captured by conventional GWAS. The reference (Ref) and alternative (Alt) alleles are separately indicated with short bars above and below in gray, which represents RNA-Seq reads aligned to the reference genome. (**D**) The transcriptome analysis enables the inference of gene co-expression networks. (**E**) Four different gene expression models during domestication, improvement, and/or adaptation in soybean are shown.

The analysis of transcriptome variation could provide valuable insights. With the use of high-density SNPs and gene expression data, we could identify loci that regulate heritable gene expression, which can be local (*Cis*) or distant (*Trans*) (**Figure 1B**). RNA-Seq not only identifies SNPs to supplement genome re-sequencing SNPs, but also reflects variations that cannot be effectively detected with low coverage next-generation based genome re-sequencing, including insertion/deletion (InDel), copy number variation (including presence-absence variation), gene fusion, epigenetic variation (methylation) (Schmitz et al., 2013), and alternative splicing (**Figure 1C**). The gene expression data also enables the inference of co-expression networks to understand gene regulation (**Figure 1D**), as well as the identification of population differentiation, revealing gene expression differences in transcriptome resulting from domestication, improvement, or geographic adaptation (**Figure 1E**).

### QTLs identification of gene expression and exon proportion

To identify the QTLs to explain as many as possible variations in expression levels, we identified both eQTLs (genetic variations associated with gene expression) and exonQTLs (genetic variations associated with coding exon proportion). The identification of exonQTLs could increase the statistical power to capture InDels, gene fusion, and alternative splicing (**Figure 1C**) via focus on exon level. For example, an exon with InDel variation can exhibit binary expression variation, i.e., with expression reads or not, while there may be no significant expression difference at the gene level. The analysis was conducted on 496 cultivars with both RNA-Seq expression data and previous genome sequencing genotypic data. 41,743 genes and 69,980 coding exons with sufficient expression data were examined using 4,331,141 imputed SNPs with a minor allele frequency (MAF) not less than 5%.

There were 44,676 eQTLs and 16,743 exonQTLs identified for 29,286 genes in total (**Figure 2A**, **Table S2, Figure S2**). The QTLs (eQTLs and/or exonQTLs) were classified as either *Cis* or *Trans* acting based on the physical distance from the leading SNP of QTL and the target gene. *Cis* are located less than the average linkage disequilibrium decay, i.e., 100 Kb (Zhou et al., 2015), while *Trans* are located longer or on different chromosomes. Permutation tests were conducted on 4,000 genes and 7,000 exons via randomly shuffling the expression data, revealing false positive rates (FDR) of ≤ 0.1% for *Cis* and ≤ 4% for *Trans* (**Figure 2A**). On average, eQTLs explained more variation of gene expression than exonQTLs explaining for reads proportion variation of exons, and *Cis* explained more than *Trans* (**Figure 2B**). There were 26,535 genes identified with *Cis*-eQTLs (N = 13,823) and/or *Trans*-eQTLs (N = 30,853) (**Figure 2C**). And exonQTLs analysis revealed *Cis*-exonQTLs (N = 5,253) and *Trans*-exonQTLs (N = 11,490) for 9,668 genes, of which 2,751 were not identified with any eQTL.

**Figure 2.**
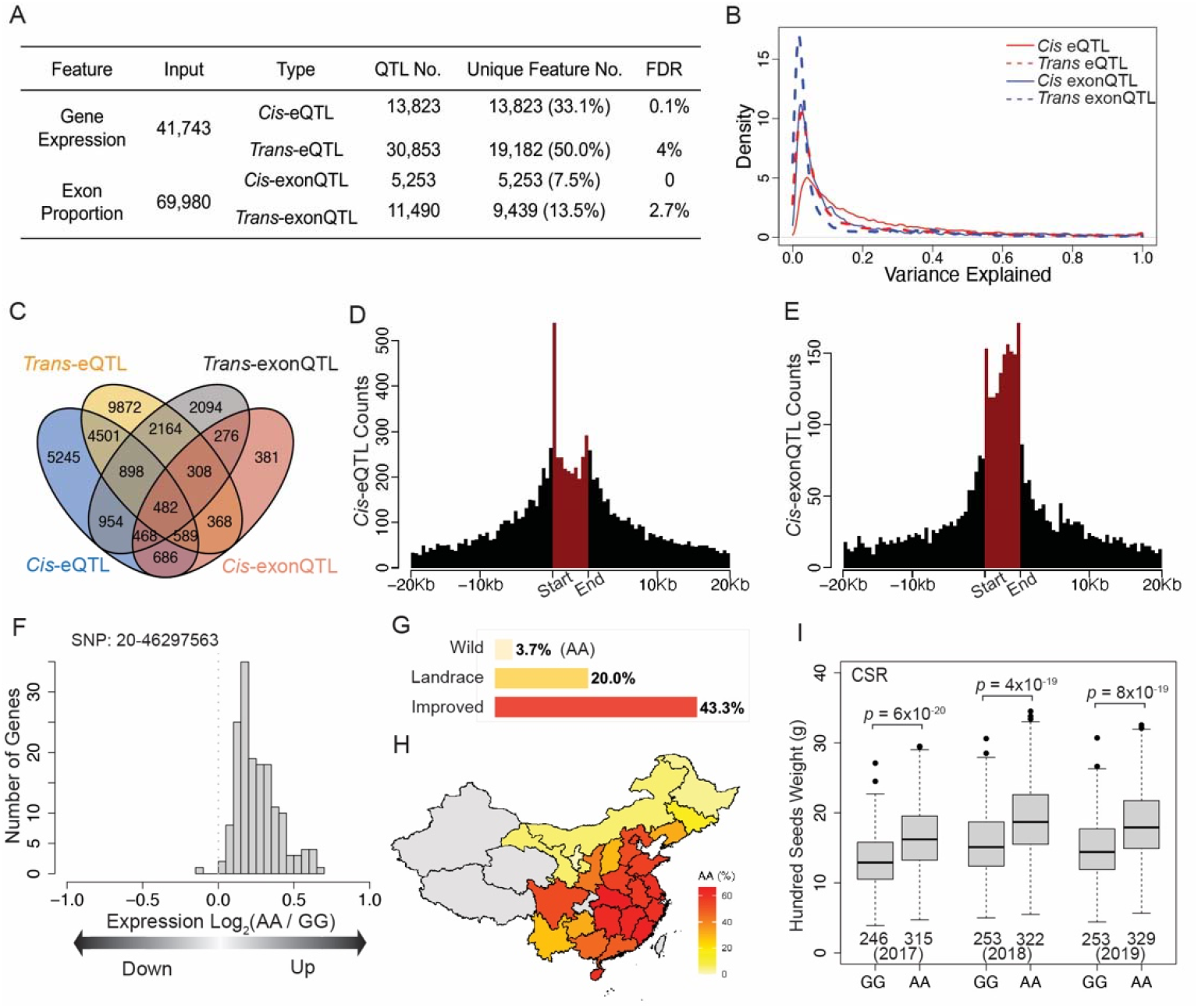
QTLs summary of gene expression and exon proportion. (**A**) Overview of gene expression QTLs (eQTLs) and exon proportion QTLs (exonQTLs). The feature represents a gene or exon, and false positive rates (FDR) was inferred from permutation of 4,000 genes and 7,000 exons for eQTLs and exonQTLs, respectively. (**B**) Variance explained by each category of QTLs. The outliers of less than 0 or higher than 1 were removed. (**C**) Overlap of regulated genes among the four QTLs categories. The number of *Cis* leading SNPs was calculated for each 200bp bin of physical distance for gene expression (**D**) and exon proportion (**E**) while +/− 20kb were plotted. The leading SNPs located within the regulated gene are summarized in red bins and scaled by their positions on the gene. (**F**) The expression of regulated genes between the two genotypes (AA and GG) of the hotspot QTN – “20-46297563”. Those genes both have “20-46297563” as the eQTL. (**G**) AA genotype frequency among populations of wild, landrace, and improved soybeans. (**H**) Percentage of AA genotype soybean cultivars in provinces of China (not all areas displayed for clarity). Provinces with fewer than 5 samples are indicated in gray. (**I**) hundred seed weight between two genotypes of the hotspot among CSR (South of China) cultivars. Three years (2017-2019) phenotype data were shown independently. Samples number with available genotype and phenotype data were indicated. The *p* values were resulted from Welch’s two-sample single tail t-test. Box plot is a visual summary of a dataset, showing the median, quartiles, and outliers. The boxes represent the middle 50% of the data and the median values as horizontal lines inside boxes. The whiskers extending from the box show the range of the data, and outliers are indicated as individual points outside the whiskers.

The *Cis*-eQTLs were defined with a physical distance of 100 Kb with the regulated genes. However, there were 2,612 (18.9%) of 13,823 *Cis*-eQTLs located within the regulated gene, and 9,488 (68.6%) had a physical distance of less than 20 Kb. For *Cis*-exonQTLs, 1,425 (27.1%) of 5,253 *Cis*-exonQTLs had leading SNP located within the regulated gene, and 3,625 (69.0%) *Cis*-exonQTLs had a physical distance less than 20 Kb. It was observed that high-density leading SNPs of *Cis*-eQTLs in the starting of genes (**Figure 2D**), and high-density leading SNPs of *Cis*-exonQTLs within the gene (**Figure 2E**). The density of both types of *Cis*-QTLs decreased with increasing physical distance with the target genes.

We identified 31 leading SNPs that *Trans*-regulated at least 33 (quantile 99.9%) genes, which were defined as quantitative trait nucleotide (QTN) hotspots (**Table S3**). Ten of these *Trans*-regulated at least 100 genes, and five of these ten QTNs were also either *Cis*-eQTLs or *Cis*-exonQTN to the gene they located in. One of these five QTNs, “20-46297563”, showed significant population differentiation, with the fixation index (F_ST_) of 0.44 (quantile 90%, 0.38) between CNR (North of China) and CSR (South of China) cultivars. The *Glyma.20G228700*, harboring “20-46297563”, had an *Arabidopsis* homology *ETHYLENE INSENSITIVE 5*, which is involved in the ethylene response pathway and showed mutant phenotype such as ethylene insensitivity, large rosettes (Olmedo et al., 2006), and delayed in bolting (Potuschak et al., 2006). The expression *Trans*-regulated genes were upregulated by the alternative genotype AA of the hotspot “20-46297563” (**Figure 2F**). The AA genotype showed increased frequency from domestication and improvement (**Figure 2G**) among the 2214 soybean accessions (Li et al., 2023b). And AA genotype had an increased frequency for geographic adaptation to high latitude (**Figure 2H**) while significantly decreased (Welch’s two-sample t-test, *p* = 3 × 10^−37^) flowering time (**Figure S3**). Additionally, the AA genotype showed significantly higher hundred seed weight among the CSR sub-population, which had the highest frequency of the AA genotype (**Figure 2I**).

### TWAS was robust with soybean seedling shoot expression data

The performance of TWAS is impacted by the source of expression data used (Li et al., 2021; Wainberg et al., 2019). To assess the performance of seeding shoot expression data, TWAS using measured gene expression was separately conducted for three qualitative traits and one quantitative trait from varying tissues and developmental stages to identify the trait-associated genes (**Figure 3**). The quantitative trait, soybean cyst nematodes (SCN) resistance, was graded on a scale of I to IV based on the resistance to race3 SCN in high latitude and long day conditions, while the seedling shoot expression data was collected in low latitude and short-day conditions. Eight known loci were identified with association signal(s) through TWAS using gene expression measurements (**Table S4**). For these signals, a 2 Mb window centered on each known gene was examined to test whether it could be uniquely identified without possible false positive (**Figure 3** and **Table 1**). Seven of those eight known loci were prioritized or uniquely identified with the known genes. The causal gene, *T* (Toda et al., 2002), for pubescence color (tawny and gray) was identified along with several associated genes surrounding it. However, these potential false-positive signals had much weaker signals and were located more than 129 Kb away. The causal *D1* and *D2* for cotyledon color (Fang et al., 2014) were identified, with a gene located 57 Kb from *D2* that may be a false positive resulting from the small sample size of green cotyledon samples (N = 14). *Rhg1* (Cook et al., 2012) and *Rhg4* (Liu et al., 2012) for SCN were known with causal multiple copy number variations (**Figure S4**). The four presumed false positives surrounding *Rhg1* and *Rhg4* for SCN were possibly caused by the shared causal multiple copy number variation, while the reason for the false positives surrounding *SNAP11* (Lakhssassi et al., 2017; Li et al., 2016) and *NSF07* (Bayless et al., 2018) remains unknown. All of the above seven known genes were either uniquely identified or prioritized. However, the causal gene *W1* (Zabala and Vodkin, 2007), responsible for determining flower color (purple and white), was expressed but was not identified as a TWAS signal. Additionally, nine surrounding genes of *W1* were identified as false positives. Collectively, the TWAS results consistent with the view that the seedling shoot gene expression data is robust for detecting trait-gene association of even non-seedling traits, but certain levels of false positive and negative rate remains a challenge for results interpretation.

**Figure 3.**
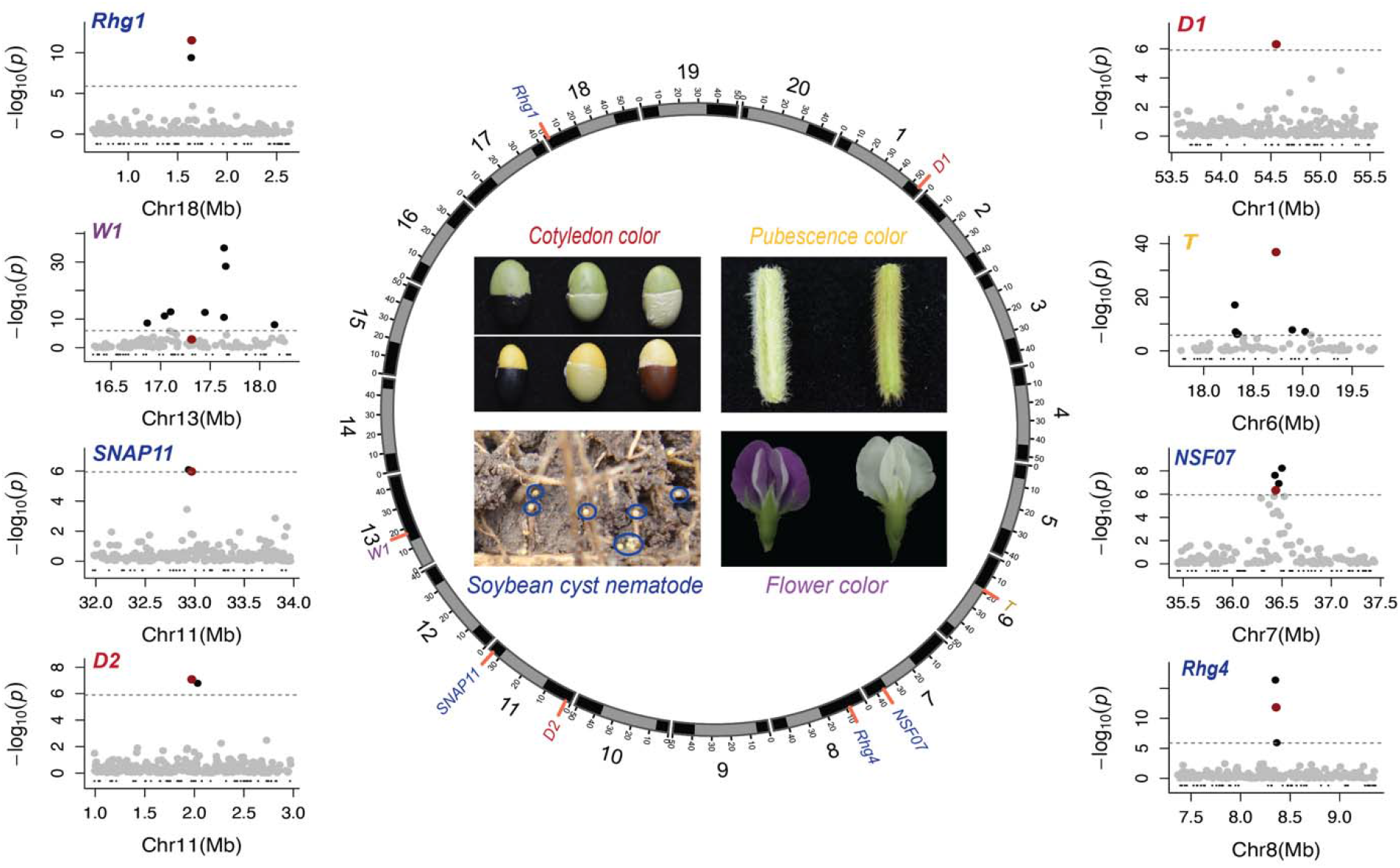
Expression of soybean seedling shoot could either uniquely identify or prioritize the known causal genes on non-seedling tissue related traits. The TWAS based on gene expression of soybean seedling shoot was conducted at four traits from four unrelated tissues: cotyledon color (green 14, yellow 480) from seed, pubescence color (tawny 253, gray 217) on the pod, soybean cyst nematode susceptibility (I 24, II 17, III 224, IV 183) from the root, and color of flower (purple 279, white 204). The identified known loci are labeled on each chromosome, with the heterochromatic regions (Song et al., 2016) indicated in grey bins. A 2 Mb window was centered on each known locus, with each dot representing an expressed gene. The names of known genes were matched with the corresponding trait names using a consistent color. Non-expressed genes were displayed in black below the −log_10_(*p*) = 0. The associated genes that passed the Bonferroni cutoff (0.05/number of expressed genes, indicated by the dashed grey line) are in black, while the known genes are in red.

**Table 1.**
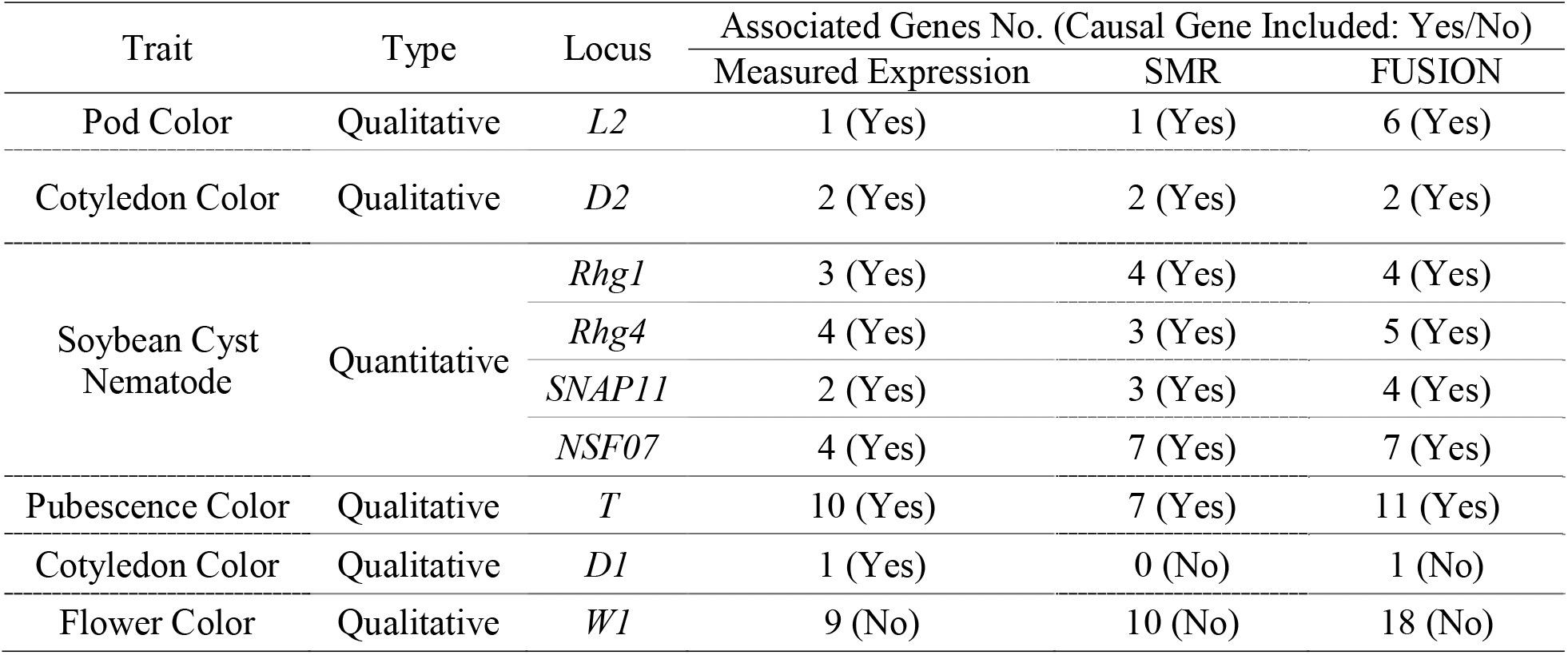
The associated genes identified by various TWAS strategies for 2Mb region centered around the causal gene.

### Pod color *L2* gene was mapped by TWAS with no effect from LD

To further evaluate the performance of TWAS, we applied it to soybean pod color trait, which has already been characterized (Bernard, 1967; Woodworth, 1923) by previous studies but the causal gene has not yet been cloned. Three distinct pod colors are controlled by two loci: black (*L1L1*, - -), brown (*l1l1*, *L2L2*), and tan (*l1l1*, *l2l2*). The GWAS and TWAS were separately performed between brown (N = 275) vs. tan (N = 68) pod color soybean cultivars to identify the *L2* locus/gene (**Figure 4A-D**). The GWAS identified a 584,234bp region (Chr03:347,610-931,843), which contained 56 annotated genes. TWAS, on the other hand, identified a single associated gene, *Glyma.03G005700*, from the 40,879 expressed soybean genes. This candidate gene located within the GWAS peak, and was 14,045 bp from the leading SNP (3-542454, *p* = 9 × 10^−25^) identified by GWAS. Two presumed functional SNPs (**Figure 4E-F**) were found that were segregating in soybean germplasm: “3-528236” induces a premature stop codon, and “3-528386” results in the only non-synonymous change between two genomes assemblies, Williams 82 (*l2*) (Schmutz et al., 2010) and W05 (*L2*) (Qi et al., 2014; Xie et al., 2019). Only two (0.9%) tan pod color samples could not be explained by one or the other of these. These two exceptions could be the result of phenotyping errors or the presence of other functional variants.

**Figure 4.**
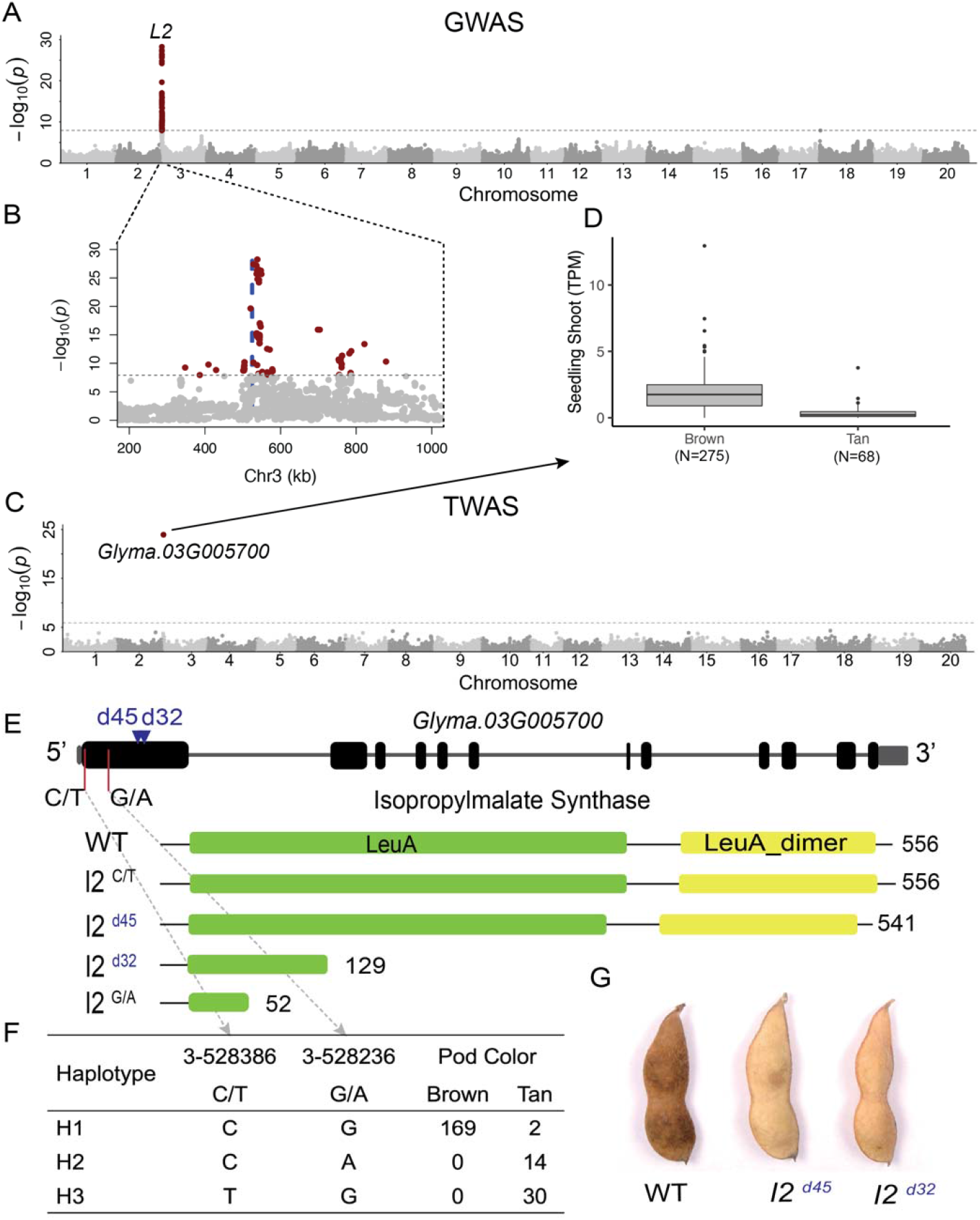
The association mapping and validation of soybean pod color gene *L2*. The GWAS (**A**) and TWAS (**B**) of soybean pod color were conducted on the same population both with compressed mixture linear model (CMLM). Each dot in (**A**) and (**B**) represents one SNP, while one gene is represented in (**C**). And associated SNPs and gene were colored in red. The blue dash line in (**B**) indicates the physical position of the single candidate gene, *Glyma.03G005700*, identified by TWAS (**C**) for the *L2* locus. The expression of the candidate gene in seedling shoot between brown and tan pod colors is shown in (**D**), normalized by transcript per million (TPM). (**E**), The gene and protein structure of *Glyma.03G005700*. The two presumed functional SNPs in the natural diversity panel were labeled in black while two CRISPR/Case9 edited events were labeled in blue. And corresponding predicted proteins were listed. (**F**), The haplotypes formed by the presumed functional SNPs (3-528236, G/A; 3-528386, C/T) and phenotype. Only the soybean accession having non-heterozygosity genotype on both two SNPs were summarized. (**G**) The pod color phenotype of the wild type (Toyomusume) and two CRISPR mutants in T_2_ generation.

To validate the candidate gene, we generated mutant alleles of the *L2* gene in the “Toyomusume”, a brown pod color soybean cultivar, using the CRISPR/Cas9 genome editing system. Two positive CRISPR events (**Figure 4E**) were identified in the first exon, one has a 45 bp deletion and another has 32 bp deletion causing a premature effect. Homozygous plants of both edits exhibited a change in pod color phenotype from the wild-type brown pod color to the mutant tan color (**Figure 4G**).

### Performance of the TWAS approaches integrating eQTLs with GWAS

An alternative approach to conducting TWAS is to integrate eQTLs with GWAS results to prioritize candidate genes. Two methods, SMR and FUSION, were applied to integrate *Cis*-eQTLs and *Cis*-exonQTLs with GWAS results for the five traits discussed above (**Table S5-S6, Figure S5**-**S6**). SMR (Zhu et al., 2016) is a Mendelian randomization method to test whether the effect size of an SNP on the phenotype is mediated by gene expression, while FUSION (Gusev et al., 2016) imputes the gene expression levels based on the genotype of GWAS panel and the expression weights of RNA-Seq panel. Both methods identified seven out of nine genes known to affect these traits, while TWAS using empirically measured expression levels identified eight of these genes (**Table 1**). Although TWAS using measured expression levels offers better performance and fewer presumed false positives, SMR and FUSION can be more widely applied as they do not require gene expression data on the subject panel for GWAS.

### Flowering time TWAS

We further applied TWAS using measured expression data to dissect the genetic architecture of flowering time (FT), which is crucial for adaptation and yield of soybean. Three kinds of expression data were separately used in the FT TWASs, including gene expression (N = 40,639), coding exons expression (N = 197,330), and coding exons proportion (N = 158,464). In total, 63 genes were associated with FT from the three TWASs (**Figure 5A**), and seven of them were known genes (**Table S7**). The genes identified via TWAS with gene expression and TWAS with exons expression exhibited a high degree of overlap, while distinct associated genes resulted from TWAS using the exons proportion (**Figure 5B**). This suggests the value of including exons proportion data to capture additional variation.

**Figure 5.**
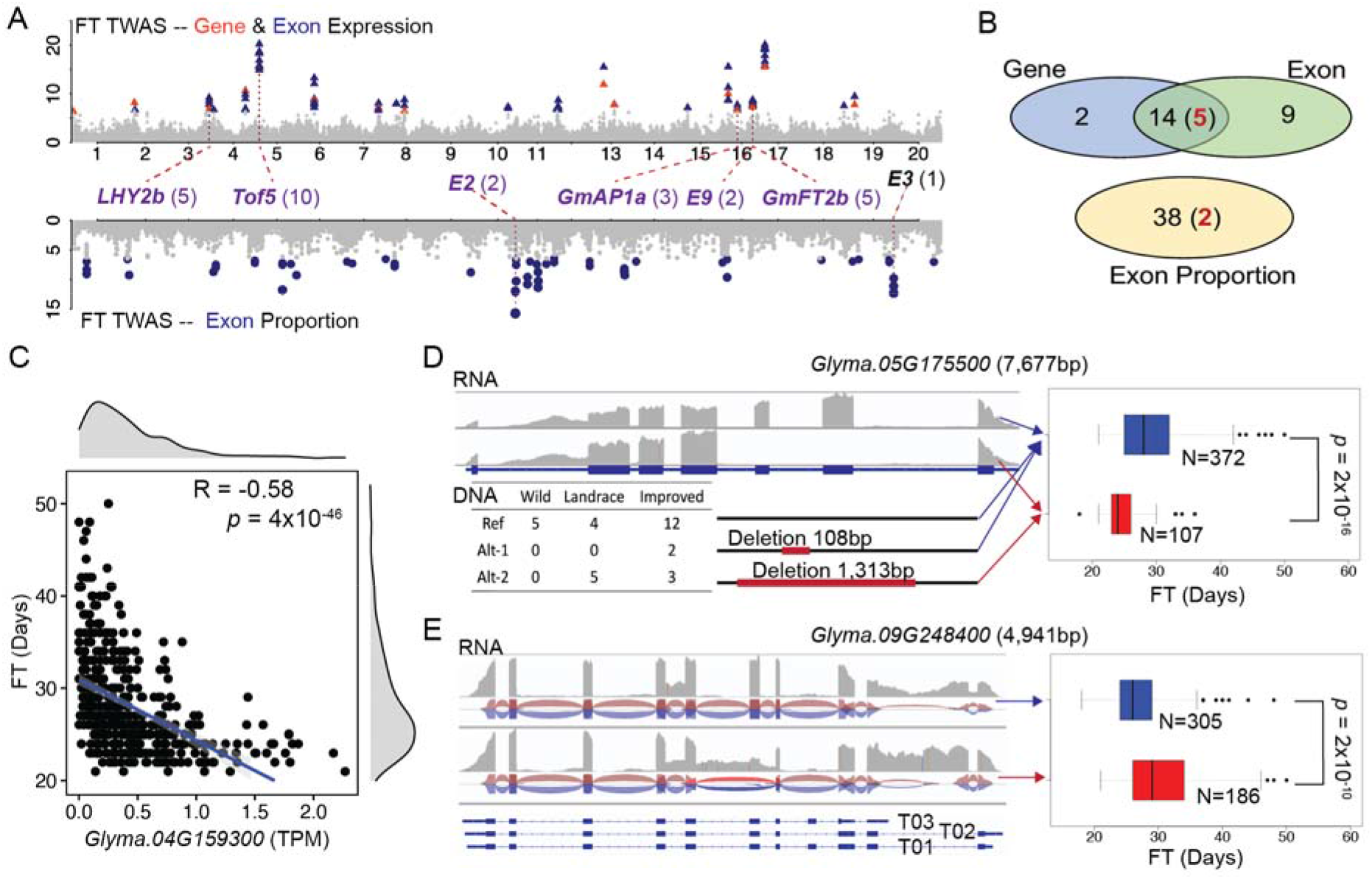
Flowering time TWAS with measured expression variations. (**A**) Three TWAS were conducted for flowering time (FT), using gene expression (red triangles), coding exon expression (blue triangles), and coding exon proportion (blue dots). The significantly associated genes/exons are highlighted in different colors, while others are in grey. The labels show the functionally known associated genes and the numbers in brackets indicate the total number of associated features (gene and exon). (**B**) Venn diagram shows the overlap of associated genes identified from each of the three different TWASs. The number of known genes, if present, is shown in brackets (red). Three novel associated genes with representation variation are shown in (**C**-**E**). (**C**) *Glyma.04G159300* had expression values significantly negatively correlated (Pearson correlation 0.58, *p* = 4 × 10^−46^) with FT. (**D**) *Glyma.05G175500* exhibited binary expression variation (expressed or not) on its two exons associating with FT. Five randomly selected samples from the top and bottom 10% expression of the two exons were pooled for visualization of alignment coverage. The deletion variation of these two exons was confirmed by a 1,313 bp deletion (Alt-2) among 31 soybean genome assemblies. The other two genotypes (Ref and Alt-1) showed no deletion of the coding sequence. The two sample categories defined by binary exon expression showed significant differences in FT (Welch’s two-sample t-test, *p* = 2 × 10^−15^). (**E**) *Glyma.09G248400* was associated with FT with alternative splicing on the sixth exon. Five randomly selected samples from the top 10% and bottom 10% expression of the sixth exon are pooled for alignment coverage and splice junction track visualization. Two sample categories defined by this alternative splicing variation showed significant FT differences (Welch’s two-sample t-test, *p* = 2 × 10^−10^).

A challenge that has been reported in TWAS is that it sometimes results in multiple hits per locus, including false positives (Wainberg et al., 2019). In this study, six out of the seven known genes identified by TWAS using various measured expression data only identified FT-related genes within 950 Kb context. *E9* (Zhao et al., 2016) and *GmFT2b* (Chen et al., 2020), located within 34 Kb of each other, were both known to regulate flowering time. The seventh known gene, *E3* (Watanabe et al., 2009), had a nearby presumed false positive gene, but this was induced by the causal gene fusion shared by both genes (**Figure S7**). Another challenge in TWAS is the potential for false positives due to co-expressed genes (Wainberg et al., 2019). To assess this, 18 modules (**Table S8**) were obtained from a co-expression network analysis. The associated genes did not show enrichment in any of those modules, suggesting that false positives resulting from correlated expression were not an issue in this case.

Three novel FT-associated genes with represented variation were further investigated. *Glyma.04G159300* showed a significantly negatively correlated gene expression (Spearman correlation 0.58, *p* = 4 × 10^−46^) with FT and is homologous to the known flowering time gene *Arabidopsis FRUITFULL* (Balanzà et al., 2014) (**Figure 5C**). The association of *Glyma.05G175500* with FT was due to the observation of expression or non-expression of its two exons. The binary expression variation was caused by variation of DNA deletion (**Figure 5D**), which was further confirmed using 31 soybean genome assemblies (Liu et al., 2020; Schmutz et al., 2010; Shen et al., 2018; Valliyodan et al., 2019; Xie et al., 2019) (**Table S9**). *Glyma.09G248400* was associated with FT through alternative splicing that involved the inclusion or exclusion of the 6th exon (**Figure 5E**), while neither RNA-Seq data nor the 31 soybean genomes revealed any InDel variations harboring the 6th exon.

We also conducted the TWAS by integrating eQTLs and exonQTLs with GWAS results on FT. The GWAS was performed with 1,715 soybean cultivars from the previous genome re-sequencing panel (Li et al., 2023b) and identified three known *FT* loci: *E1* (Xia et al., 2012), *E2* (Tsubokura et al., 2014), and *GmPRR3b* (Li et al., 2020a) (**Figure S8**). *E2* was the only gene identified with *Cis*-eQTL or exonQTL, its presumed stop-gained causal SNP were both identified as the leading SNP in eQTL and FT GWAS (**Figure S9**). *E2* was identified as FT-associated (**Figure S8**, **Table S5-S6**) by SMR, but not FUSION, without any other signals surrounding a 200 Kb region centered on the gene, which is twice the size of the average LD decay in soybean (Zhou et al., 2015). On the other hand, seven of those 63 FT-associated genes were identified with *Cis-* QTL only, 16 were identified with *Trans-* QTL(s) only, and 36 were identified with *Cis-* and *Trans-* QTLs. Four genes, including two known genes, were identified with no eQTL or exonQTL.

### Transcriptome insight into soybean domestication, improvement, and adaptation

Population differentiation from genotype analysis can reveal genetic loci related to evolution, improvement, and adaptation. In this study, we identified the differentially expressed genes (DEGs) (**Table S10**) between wild ancestors and landrace of soybean for domestication-related genes (N = 2,673); landrace and improved cultivars for improvement-related genes (N = 429); among three cultivars geographic sub-populations for two adaptation-related genes sets (N = 2,921) (**Figure 6A-B**). The “ADP binding” GO term (GO:0043531) was enriched in each of the four DEG sets (**Table S11**). FT, a trait undergoing selection during domestication, improvement, and adaptation, had 37 of its 63 associated genes differentially expressed in at least one of the four comparisons, with four known FT-related genes found in at least one of DEG sets. One of these is *Tof5*. *Tof5* is a gene that has been reported to have undergone domestication selection and is known to exhibit polymorphism only in high latitudes, such as northern China (Dong et al., 2022). The differential expression of *Tof5* between wild and landrace soybeans (**Figure 6C**) consistent with the view that this gene is likely under selection during the domestication process. In addition, *Tof5* shows a gradient expression pattern, which may be influenced by its regulation by both *Cis*- and *Trans*-eQTLs. This gradient expression is correlated with both originated latitude (Spearman correlation 0.57, *p* = 2 × 10^−40^) and FT (Spearman correlation −0.66, *p* = 1 × 10^−62^) in soybean cultivars (**Figure 6D**). Neither *Tof5*, nor any of the *Trans-*regulators of *Tof5* identified via eQTL analysis, were identified in the conventional GWAS for FT. However, TWAS using the expression level of *Tof5*, which integrates the effects of one *Cis*-eQTL, six *Trans*-eQTLs and one *Trans*-exonQTL was able to accurately link this gene to a role in flowering time (**Figure S10**).

**Figure 6.**
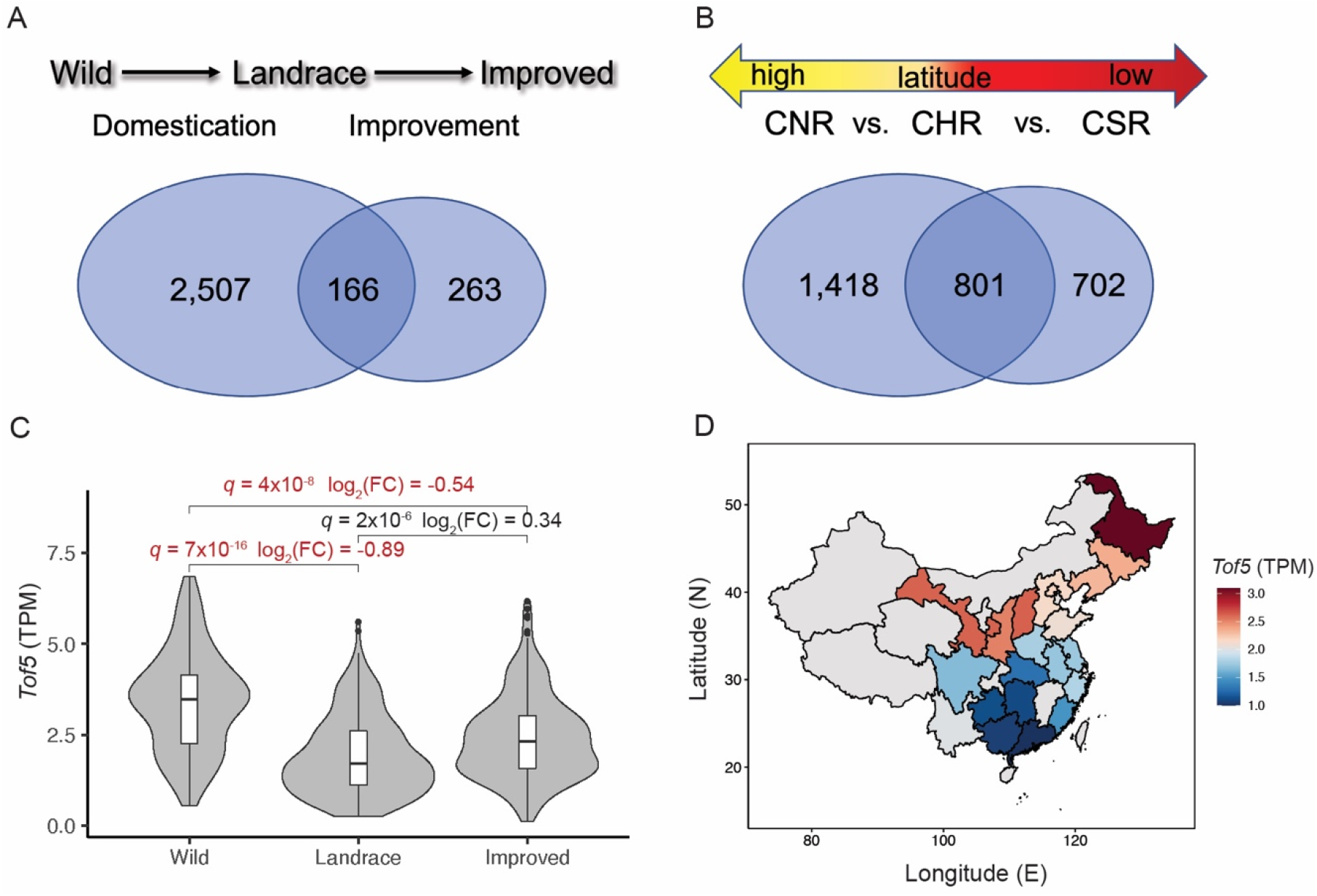
Differential gene expressions between sub-populations. (**A**-**B**) Venn diagrams showing the overlap between the different sets of differentially expressed genes (DEGs). (**C**) Gene expression of *Tof5* across different groups. *q* value (adjusted *p*) and fold change (FC) information were obtained from DEGs analysis, the comparisons passed the cutoff (*q* <0.01 and |log_2_(FC)| >0.5) were highlighted in red. (**D**) The average expression of *Tof5* in RNA-Seq data from cultivars based on their origin from different provinces in China (not all areas displayed for clarity). Provinces with fewer than 10 samples are represented in gray.

## Discussion

In this study, self-pollinated soybean has been used as a model for slow LD decay plants to demonstrate how transcriptome data could facilitate gene mapping. We analyzed the first transcriptome variation of a soybean natural population and bridged the gap between genome variation and transcriptome variation by identifying genetic loci (eQTLs and exonQTLs) that regulate expression variation for 29,286 genes. TWAS, using different approaches, demonstrated the ability to provide gene-level association mapping for different traits in soybean. This was not only evident in cases involving known genes, but also in identifying and functionally validating a gene, *L2*, first described based upon phenotype exactly one hundred years ago (Woodworth, 1923).

Previous studies (Li et al., 2021; Sun et al., 2023; Zheng et al., 2020) have demonstrated that TWAS is robust in detecting the causal gene using expression data obtained from tissues that are not directly related to the trait of interest. However, others have raised concerns that, given the dynamic nature of the transcriptome across tissues, developmental stages, and in response to environmental factors, selecting the appropriate samples may be critical for successful TWAS (Finucane et al., 2018; GTEx Consortium, 2017; Wainberg et al., 2019). Our study found that seedling shoot expression data was effective in detecting true positive genes for five out of six traits we evaluated, including traits observed in different tissues, developmental stages, and/or environmental conditions. Notably, we demonstrated that seedling shoot expression data was also effective in TWAS analysis for resistance of SCN, which is microscopic roundworms that infect the roots. The robustness of the TWAS results observed here suggests that the same expression dataset could well be combined with phenotyping data collected in a wide range of environmental conditions to provide insight into the genes controlling variation in phenotypic responses to those same environmental factors.

The access to technology for profiling alternative splicing on a genome-wide scale has enabled the discovery and quantification of significant variation in alternative splicing within and across populations (Gan et al., 2011; Wang et al., 2008). Alternative splicing QTLs (sQTLs) have been identified in both animal and plant systems and naturally occurring genetically controlled variation in alternative splicing has been linked to variation in phenotypes in both humans (Qi et al., 2022) and plants (Chen et al., 2018; Sun et al., 2023). However, current approaches for integrating analysis of alternative splicing and phenotypic variation in plants typically have depended on colocalization of GWAS hits and sQTLs, rather than employing genome-wide patterns of alternative splicing as explanatory variables to explain variation in plant traits. Here we introduced a new feature called “exon proportion” which captures variation resulting from both alternative splicing and structural variation, another type of functional genetic variation which is abundant in plant genomes (Hirsch et al., 2014; Jayakodi et al., 2020; Liu et al., 2020; Wang et al., 2018) but frequently missed in conventional GWAS. Exon proportion features calculated from RNA-Seq data of a large soybean diversity panel were employed to identify the regulation of genetic loci (exonQTLs) and to perform association studies with phenotype variation (TWAS). Our exonQTL analysis identified *Cis-* or *Trans-* exonQTLs for 2,751 genes that had no eQTLs. Using the feature “exon proportion” in TWAS leds to the identification of unique genes, including genes previously demonstrated to be true positives, which were not identified using conventional TWAS with gene and exon expression (**Figure 5A-B**). Our analysis revealed the impact of different structural variations and alternative splicing on gene expression related to flowering time. Specifically, we identified a naturally occurring gene fusion effective the well-known *E3* (Watanabe et al., 2009), an InDel in *Glyma.05G175500*, and variation of alternative splicing at the *Glyma.09G248400* locus associated with flowering time (**Figure 5C-E**).

It has been hypothesized (Li et al., 2021) that the frequent observation that TWAS identified additional true positive genes not captured by GWAS in the same populations may be due to the presence of unrepresented *Cis*-eQTL or epigenetic variation or the result of *Trans*-eQTL. In the latter case, gene expression can also be influenced by the presence of multiple regulators, each of which may have a small effect size. In combination, their effects can amplify the expression level of the regulated genes. In the case of the flowering time regulatory *Tof5* (**Figure S10**), we see a result extremely consistent with the *Trans*-eQTLs model, with the model small effect *Trans-*eQTLs regulating the expression of *Tof5* and the expression of *Tof5* in turn linked to variation in flowering time while the *Trans*-eQTLs individually do not exhibit statistically significant links to flowering time GWAS in this study. One implication of this *Trans*-eQTLs model for why TWAS identifies unique genes is that it may help to identify important regulators where little or no functional variation is currently segregating, such as genes that were targets of selection during domestication.

## Materials and Methods

### Soybean germplasm and phentoyping

This study employed a previously described soybean diversity panel, consisting of 2,214 re-sequenced lines (Li et al., 2023b) (**Table S12**). Measurements of flowering time and hundred seed weight were collected from field studies conducted in Nanchang (N 28°31′, E 116°1′) in 2017, 2018 and 2019. A subset of the FT data from these field studies has been described in a previous publication (Li et al., 2023a). Each soybean accession was planted as a 1.8 × 0.8 m plot with two rows, with 10 cm between seedlings for each row. FT and hundred seed weight data were averaged from individuals of those adjacent two rows. FT was scored as the number of days from the emergence of the cotyledons to the first day when open flowers were present on at least 50% of the plants within the row. Values for each accession were averaged across all three years.

The resistance to race3 SCN was assessed in Harbin (N 45°45′, E 126°41′) during 2017 and 2018 using a standardized I-IV scale, based on the female index (FI): I (highly resistant, FI <10%), II (moderately resistant, 10%≤FI<30%), III (moderately susceptible, 30%≤FI<60%), and IV (highly susceptible, FI≥60%). FI was calculated for each plant by the number of female cyst nematodes on a given plant to the average number on susceptible plants. A previously described scoring metric (Li et al., 2016) was used, which planted each genotype with three replications, and creating one 1.5 × 0.65 m row for each replication, with a 5 cm spacing between seedlings. Five individuals in the center of row were graded at thirty days after emergence. Data on qualitative traits, including pod color, cotyledon color, flower color, and pubescence color, was obtained from the records of the Chinese National Soybean GeneBank. As a quality control measure, pod color was re-phenotyped in Nanchang in 2018. Only data for soybean accessions with consistently recorded pod colors were used in this study.

### Gene expression quantification

RNA-Seq data was generated for a set of 622 soybean accessions, including 560 accessions shared with the genome re-sequenced population described above (**Table S1**). This RNA-Seq population included 70 wild, 304 landrace, and 248 improved soybean lines from around the globe. The seeds of each accession were sown in field with an average light time of 11 hours 20 minutes and a temperature of 22°C in Sanya (N 18°23′, E 109°9′). Tissue samples consisting of all tissue above the cotyledon node for 2–3 individual plants were collected at the V2 developmental stage for each accession. To reduce the confounding effects of diurnal cycles on variation in gene expression, all tissue samples were collected between 9 and 11 a.m.

Total RNA was extracted using RNA Easy Fast Plant Tissue Kits (Tiangen Biotech, China). RNA integrity was assessed using the RNA Nano 6000 Assay Kit of the Bioanalyzer 2100 system. mRNA was purified from total RNA by using poly-T oligo-attached magnetic beads. RNA-Seq libraries were constructed using the NEBNext^®^ UltraTM RNA Library Prep Kit. cDNA fragments between 370 bp and 420 bp in length were selected, purified, and amplified with AMPure XP system (Beckman Coulter, Beverly, USA). The resulting libraries were sequenced using the Illumina Novaseq 6000 with 2×150 base pair sequence reads.

Trimmomatic (Bolger et al., 2014) (V0.36) was used to quality control and remove low-quality bases or reads with parameter settings (LEADING:15 TRAILING:15 SLIDINGWINDOW:4:20 MINLEN:100). The quality filtered and trimmed pair-end reads were aligned to the soybean Williams 82 V2 reference genome (Schmutz et al., 2010) using STAR (Dobin et al., 2013) (V2.7.1a) (–outFilterMultimapNmax 10 –alignIntronMax 50000). Only uniquely aligned reads were used for subsequent analyses. Reads counts were separately obtained for entire genes and individual coding exons using featureCounts (Liao et al., 2014) (-s 0 -p -B -C) while allowing reads to be assigned multiple matched meta-features. The expression values of gene and coding exons were normalized with transcripts per kilobase million (TPM) method separately.

### Genotyping

Uniquely aligned reads were used to discover and score SNPs using GATK (V4.1.5) and the parameters described in “RNAseq short variant discovery” workflow (Van der Auwera et al., 2013). 480,785 raw SNPs set was obtained after discarding those SNPs which were not bi-allelic, genotyped in <50% of samples, or where the minor allele was detected in less than10 samples. A high quality SNP set was generated from the raw SNP set by retaining only the subset of markers where: 1) MAF was not less than 5%; 2) the minor allele was scored as homozygous at least 10 samples; 3) <20% of the population was scored as heterozygous; 4) The analysis retained filtered SNPs but removed those located on scaffolds. The high-quality chromosome-based SNPs were retained for subsequent analyses. The new SNP set derived from RNA-Seq and previous SNP set derived from genome re-sequencing (Li et al., 2023b) (N = 8,785,134) were separately imputed using beagle (Browning and Browning, 2016; Browning and Browning, 2007) (V5.0). After imputation, records for individuals present in both the RNA-Seq set and the genome re-sequencing set were merged, with the genotype observed in the RNA-Seq set retained when the two records disagreed. The functional effect for each individual SNP in the dataset was predicted using SNPEff (Cingolani et al., 2012) (V4.3t).

### eQTLs and exonQTLs

Mapping of quantitative trait loci controlling variation in both gene expression and exon proportion was conducted using data for a set of 496 domesticated soybean cultivars (273 landrace and 223 improved), for which both expression data and whole genome re-sequencing genotyping was available. For eQTL analysis, informative genes, defined as those with an average expression value >0.1, were normalized across samples using the qqnorm function implemented in R language (V4.1.2) with a normal quantile transformation. Exon proportion values, which is the ratio of the number of reads mapping to a given exon to the total number of reads mapping to gene, were calculated for genes with at least two exons. The exons were required to have an average of > 20 aligned reads among the 496 cultivars. To reduce the redundancy, annotated exons with identical start and end positions were removed. In addition, in any case where two exons exhibited a spearman correlation coefficient >0.6, the second exon (e.g. the one more proximal to downstream of the gene) was dropped from the analysis. Exon proportion values for the exons that passed this filter were normalized using the normal quantile transformation prior to exonQTL analysis.

GWAS for each gene and exon proportion were conducted using BLINK (Huang et al., 2019) (V0.01) to identify associated SNPs. The merged and imputed RNA-Seq based high-quality SNPs and previous re-sequencing based SNPs were filtered with MAF ≥ 5%, the retained 4,331,141 SNPs were used for GWAS. Population structure was controlled for using the first three components of genotypic variation as determined by a principal component analysis (PCA) conducted in PLINK (Purcell et al., 2007) (V1.9).

Based on the rate of LD decay observed in soybean (Zhou et al., 2015), QTLs of gene expression and exon proportion were defined as *Cis* when the distance between the leading SNP of the QTL and the gene was <100 Kb and defined as in *Trans* in all other cases. Because the degrees of multiple testing present for *Cis*- and *Trans*-QTLs were substantially different, a *p* value cutoff of 1 × 10^−6^ was applied for *Cis*-QTLs, while a Bonferroni correction at level α = 0.05 *p* value cutoff, i.e., 1.2 × 10^−8^, was required for a *Trans* to be considered significant. Associated SNPs identified in GWAS analyses were filtered before defining the QTLs. Since each associated SNP resulted from BLINK should, in principle, represent an independent QTL. In cases where >50 SNPs with *p* values less than 1 × 10^−6^ were associated with a single gene (N = 285; 0.7% input genes) or exon (N = 2,040; 2.9% input exons), these genes and exons were considered noise results and dropped from the analysis. And SNPs either with a LD R^2^ >0.4 or distance <100 Kb were further merged with the leading SNP. The associated SNPs were also required to have a heterozygosity rate ≤ 20% and at least 5 minor allele homozygous samples. Variance explained by each leading associated SNP was calculated using SNPs within its centered 200 Kb via PLINK (Purcell et al., 2007) and GCTA (Yang et al., 2011) (V1.94.1).

Permutation tests were used to evaluate the false positive rate of the eQTLs and exonQTLs separately. 4000 genes and 7000 exons were randomly selected to randomly shuffle the sample and expression values for the test, and the same analysis criteria described above were used to identify QTLs that should not be identified, creating an estimate of the false discovery rate.

### QTN hotspots

The leading SNP, i.e., QTN, from each *Trans*-eQTLs and exonQTLs were counted after removing of redundant QTN-Gene (N = 33) resulting from *Trans*-exonQTLs results. Weir and Cockerham’s F_ST_ (Weir and Cockerham, 1984) was calculated using VCFtools (Danecek et al., 2011) (V0.1.17) for pairwise sub-populations on each of those QTN also all other SNPs used for eQTLs analysis.

### GWAS of agronomic traits

GWAS for both qualitative and quantitative agronomic traits were conducted using the compressed mixture linear model (CMLM) (Zhang et al., 2010) implemented in GAPIT (Lipka et al., 2012) (V3.1). For each analysis, the SNP set was independently filtered to retain only SNPs with a MAF of >5% among samples with recorded trait values, and for each analysis population structure was controlled for using the first three components of the genotypes calculated using a PCA of available samples. A Bonferroni correction of level α = 0.05, i.e., 0.05/SNPs number, was used to define the associated SNPs.

### Different TWAS strategies

Two strategies to TWAS were employed in this study. The first of these two was to perform association tests between directly measured expression levels – here quantified in three ways as normalized gene expression, normalized coding exons expression, and exon proportion – and phenotypic values (“TWAS with measured expression data”). All three types of quantified expression data were converted to the 0-2 range required by GAPIT using the method previously described (Li et al., 2021), which converts bottom 5% ( ) and top 5% ( ) values to 0 and 2 separately, and then linearly transformed the other values from 0 to 2 (**equation (1)**). After conversion, the CMLM method as implemented in GAPIT (Lipka et al., 2012) (V3.1) used for GWAS of agronomic traits was also employed for TWAS using measured expression data. The Bonferroni correction at level α = 0.05, i.e., 0.05/number of genes or gene features, was used to define the associated genes or gene features.

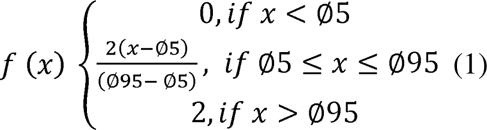

The second strategy is to integrate eQTLs into GWAS to prioritize the causal genes. Two methods, FUSION (Gusev et al., 2016) and SMR (Zhu et al., 2016) (V1.3.0) for this integration, were employed in this study. Only *Cis*-eQTLs and *Cis*-exonQTLs were integrated, and centered-200Kb SNPs of each leading associated SNP and the univariate linear mixed model implemented in GEMMA (Zhou and Stephens, 2012) (V0.98.1) were used for expression weights (–models top1, lasso, enet) required by FUSION and summary-level data required by SMR, with other default parameters. And the required GWASs for those tools were also conducted with GEMMA using the univariate linear mixed model. The Bonferroni correction at level α = 0.05 was used for the *p* value cutoff for both methods.

### Variations confirmation on soybean pan-genome

Structural variants inferred from RNA-Seq alignments were validated using independent genome assemblies for 31 soybean accession (Liu et al., 2020; Schmutz et al., 2010; Shen et al., 2018; Valliyodan et al., 2019; Xie et al., 2019). In each case, the region where a structural variant was identified from RNA-Seq alignments was manually quality controlled using IGV (Robinson et al., 2011) (V2.11.9), and a region beginning 100 bp upstream of the start of the variant and extended 100 bp downstream of the variant was extracted from the Williams 82 (V2) reference genome. This extracted region was used as a query for a blastn (Camacho et al., 2009) (V2.13.0) search against each of other 30 soybean reference genomes to identify the location and size of an equivalent genomic region. These corresponding regions were extracted, and subsequent multiple sequence alignment of genomic DNA sequences was constructed using MUSCLE (Edgar, 2004) implemented in EMBL-EBI (Madeira et al., 2022).

### Genome editing of *L2*

To construct the plant transformation plasmids, the sequence and information for the *L2* gene (*Glyma.06G207800*) was downloaded from the Phytozome database (https://phytozome-next.jgi.doe.gov) and an online tool (http://www.genome.arizona.edu/crispr/CRISPRsearch.html) (Xie et al., 2014) used to design two optimal sgRNA sequences for the *L2* gene. Two sgRNAs named L2-SP1 (5’-CTGTCATTGCGGCCCTATGT-3’) and L2-SP2 (5’-GGACATAACGAGAGCCTGGG-3’) were designed (**Figure S11**). The two sgRNAs were cloned into the pHSE401 plasmid vector to construct the CRISPR/Cas9 expression vector. The *Agrobacterium* strain EHA105 was transformed with this expression vector using the electroporation method. The cotyledon-node method (Paz et al., 2006) was employed to culture tissues and transform the brown pod color soybean cultivar Toyomusume. Homozygous mutants were obtained from the self-pollination of candidate mutants, and the edited sites were verified by Sanger sequencing with the primers in **Table S13**.

### Co-expression network analysis

A weighted co-expression network was inferred from all RNA-Seq sample expression data using the R package WGCNA (Langfelder and Horvath, 2008) (V1.70.3). Genes with average reads numbers greater than 20 were log-transformed using log_2_(TPM+1) and used for network construction. Outlier samples were removed (cutHeight = 130, minSize = 10) prior to network construction. Subsequent parameters (power = 12, minModuleSize =100, cutHeight = 0.2) were adjusted to generate co-expressed modules.

### Differential expression genes analysis

Differential gene expression analysis was performed using the R package DESeq2 (Love et al., 2014) (V1.34.0). Prior to analysis, outliers identified during co-expression network analysis were removed. Genes having average aligned reads exceeding 20 among subject samples were considered for the input of analysis, along with the raw reads numbers. The accessions in each group were treated as replicates, and a generalized linear model was used for the analysis. The DEGs were defined as those with an adjusted *p* value (*q* value) of less than 0.01 and an absolute value of log_2_(FoldChange) greater than 0.5.

### Functional enrichment analyses

Functional enrichment analysis was performed using the R package clusterProfiler (Wu et al., 2021) (V4.0) for the enrichment of gene ontology (GO) terms and the co-expression gene modules. The significance of each enriched term/module was assessed using a Benjamini-Hochberg correction for multiple testing. Terms/modules with an adjusted *p* value (*q* value) less than 0.01 were defined as significantly enriched.

### Data availability

The raw RNA-Seq sequencing data generated in this study have been deposited into the Genome Sequence Archive (GSA) database in BIG Data Center (https://bigd.big.ac.cn/gsa/index.jsp) as accession number CRA009979. The processed reads counts for gene and coding exon, expression weights required by FUSION, summary level data required by SMR, and pod color TWAS with three methods are available at figshare (https://figshare.com/s/811b3ef0dfc6cba0a167).

## Author contributions

Y.H.L., J.C.S, L.J.Q., and D.L. conceived the research, D.L. performed the data analysis, X.L., Y.S., B.L., Q.W., H.Z., and D.L. performed gene validation. Y.F.L. prepared samples for RNA-Seq. Y.T., C.Z., H.H., H.G., J.W. and R.W. collected phenotyping data. J.Y and P.S.S contributed ideas and discussions on the analysis. Y.H.L., J.C.S, and L.J.Q. supervised the research. D.L., J.C.S., and Y.H.L. wrote the manuscript. All authors reviewed and approved the final manuscript.

## Acknowledgements

We would like to thank Dr. Zhiwu Zhang (Washington State University) and Dr. Weifeng Zhang (Tsinghua University) for fruitful discussions, and thank the Core Facility Platform, Institute of Crop Sciences, Chinese Academy of Agriculture Sciences (CAAS), for assistance with partial sequencing. This work was supported by the National Key Research and Development Program of China (2021YFD1201601), National Natural Science Foundation of China (32201759, U22A20473), the earmarked fund for CARS (CARS-04-PS01), and the Agricultural Science and Technology Innovation Program (ASTIP) (CAAS-ZDRW202109).

## Supporting information

**Table S1. Material information and sequencing data summary.**

**Table S2. eQTLs and exonQTLs.**

**Table S3. The 31 hotspots of eQTLs and exonQTLs.**

**Table S4. Associated genes resulted from TWAS using measured gene expression for the four known traits.**

**Table S5. TWAS SMR associated results.**

**Table S6. TWAS FUSION associated results.**

**Table S7. Results of flowering time TWASs with measured expression.**

**Table S8. Modules of co-expression network.**

**Table S9. InDel variation of *Glyma.05G175500* in 31 soybean assemblies.**

**Table S10. DEGs lists of the four comparisons.**

**Table S11. GO enrichment of DEGs lists.**

**Table S12. Phenotype data.**

**Table S13. Primers for verification of *l2* mutants editing sites.**

**Figure S1. Represented samples were selected from the genome re-sequencing panel for RNA-Seq.**

**Figure S2. Genome-wide distribution of different gene sets.**

**Figure S3. Flowering time between genotypes of the hotspot “20-46297563”.**

**Figure S4. Gene expression of *Rhg1* and *Rhg4*.**

**Figure S5. TWAS SMR for each of the five known traits.**

**Figure S6. TWAS FUSION for each of the five known traits.**

**Figure S7. Gene fusion of *E3* revealed with RNA-Seq data.**

**Figure S8. Association analyses of flowering time.**

**Figure S9. Gene expression and flowering time for genotypes of the casual variant of *E2*.**

**Figure S10. Gene expression and flowering time for the different pseudo-haplotypes of eQTLs and exonQTL of *Tof5*.**

**Figure S11. The gRNAs for genome editing of *L2*.**

