## Supplementary material for "Transcriptome brings variations of gene expression, alternative splicing, and structural variations into gene-scale trait dissection in soybean": Figure S

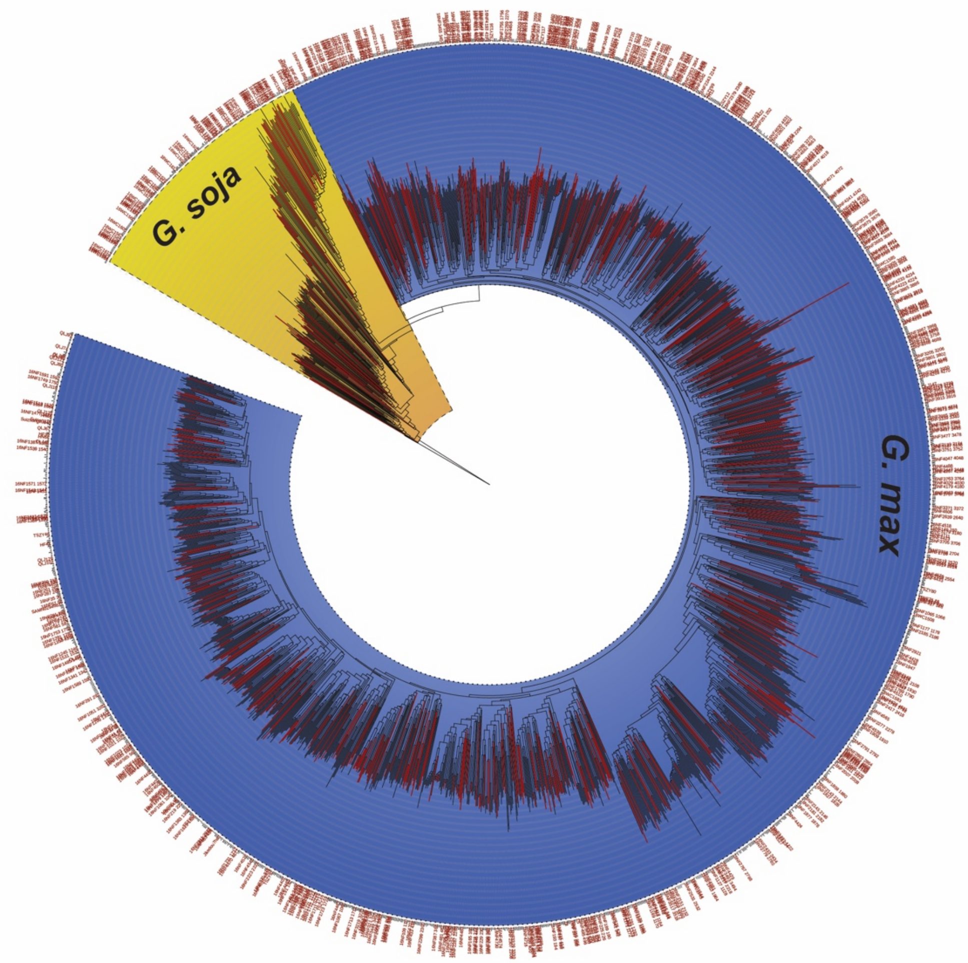


**Figure S1. Represented samples were selected from the genome re-sequencing panel for RNA-Seq.**

Phylogenetic tree (Li et al., 2023b) of the genome re-sequencing panel. The wild soybean (*G. soja*) and cultivars (*G. max*) were highlighted. There are 560 soybean accessions selected to represent the panel for RNA-Seq, those RNA-Seq accessions have labels and branches highlighted in red.

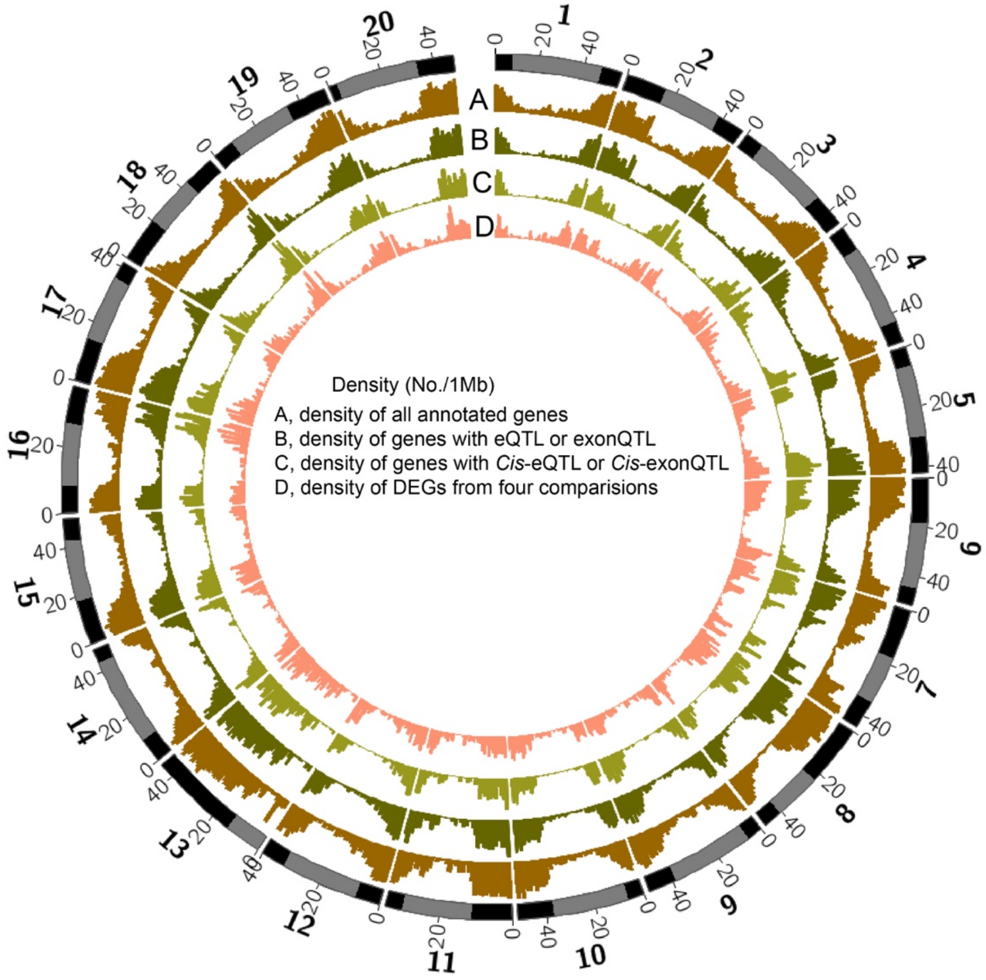


**Figure S2. Genome-wide distribution of different gene sets.**

The gene number per 1 Mb for four different gene sets: (**A**) All annotated gene in Wm82 V2; (**B**) Genes were either identified with eQTL or exonQTL; (**C**) Genes were either identified with *Cis*-eQTL or *Cis*-exonQTL; (**D**) Genes differently expressed in at least one of four DEGs analyses. The chromosome heterochromatic regions (Song et al., 2016) were indicated in grey, while other regions are shown in black.

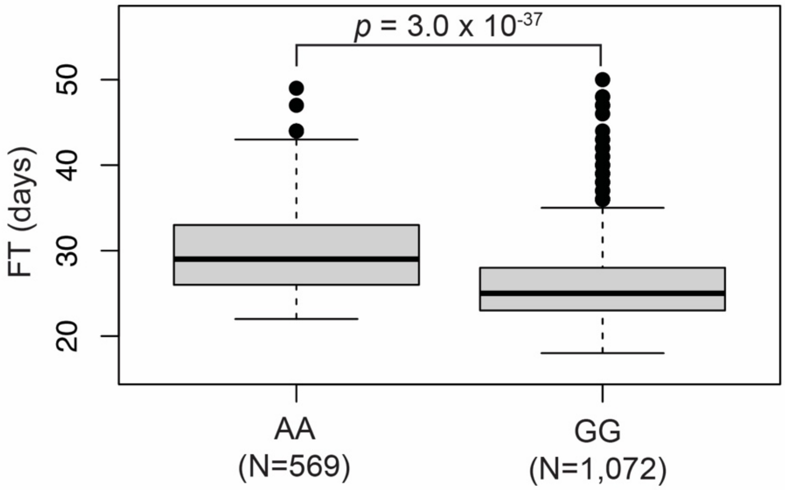


**Figure S3. Flowering time between genotypes of the hotspot "20-46297563".**

The soybean cultivars studied were cultivars from a genome re-sequencing 2,214 panel having flowering time (FT) information and genotyping at "20-46297563". $p = 3 \times 10^{-37}$ was resulted from Welch's two-sample t-test. The studied soybean cultivars came from the genome re-sequencing panel that included flowering time (FT) information and genotyping at SNP of "20-46297563".

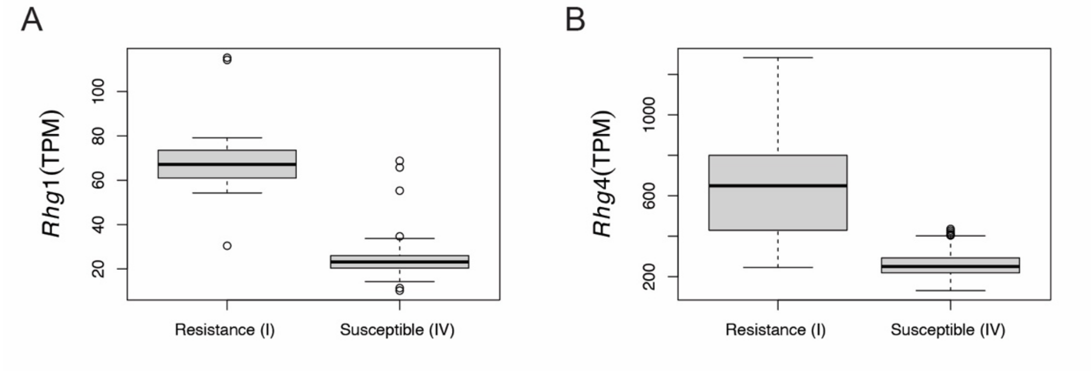


**Figure S4. Gene expression of *Rhg1* and *Rhg4*.**

Gene expression of *Rhg1* (A) and *Rhg4* (B) for different subset samples. After removing outlier samples with top 2% *Rgh1* expression value, the soybean samples were categorized based on their resistance grade to soybean cyst nematodes (SCN): I (N = 22), II (N = 16), III (N = 220), and IV (N = 181). Grade I represented the highest level of resistance were reported with high number copy of *Rhg1* (Cook et al., 2012) and *Rhg4* (Liu et al., 2012). The labeled number on the box are the median expression for corresponding category.

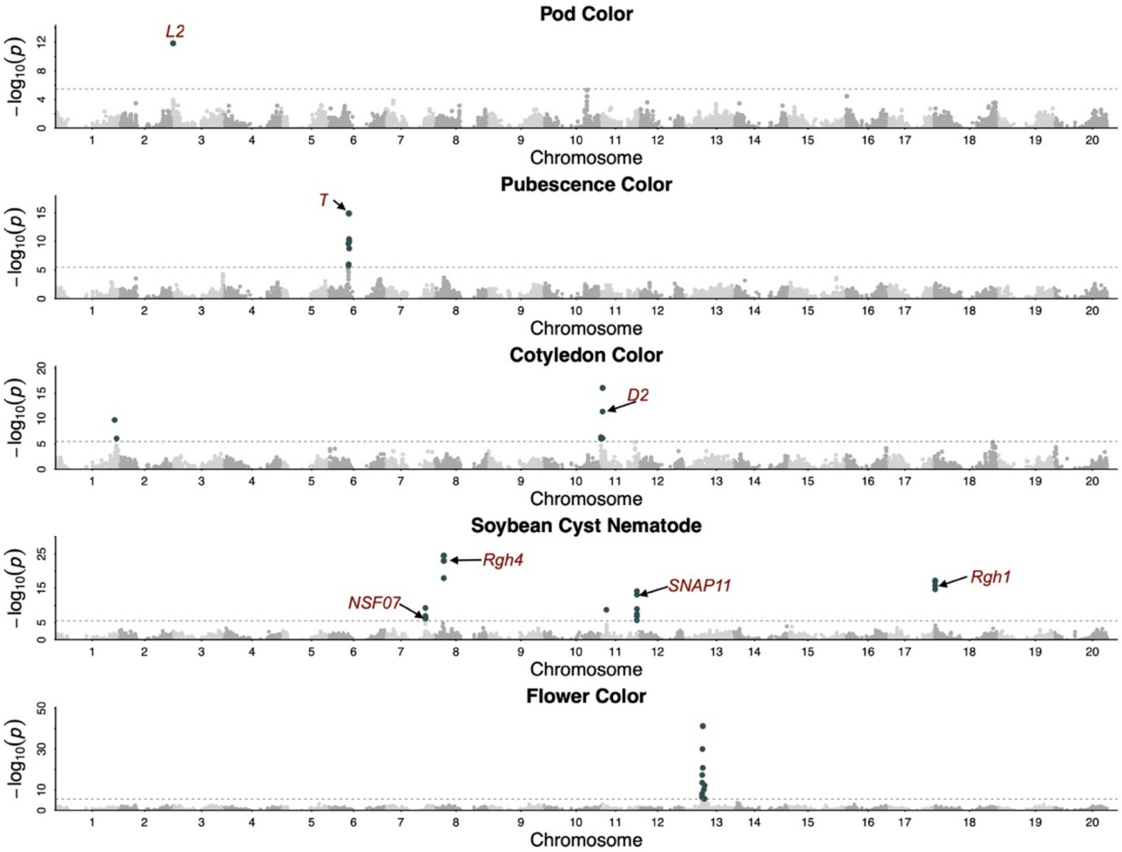


**Figure S5. TWAS SMR for each of the five known traits.**

Manhattan plots of each TWAS SMR for corresponding traits. Dashed line indicates the association cutoff. Each dot on the plot represents a feature, i.e., gene with *Cis*-eQTL or exon with *Cis*-exonQTL. The associated features that were found to be significant are highlighted in black, while non-significant ones are shown in grey. When known genes were identified as significantly associated, they were labeled with their most significant feature.

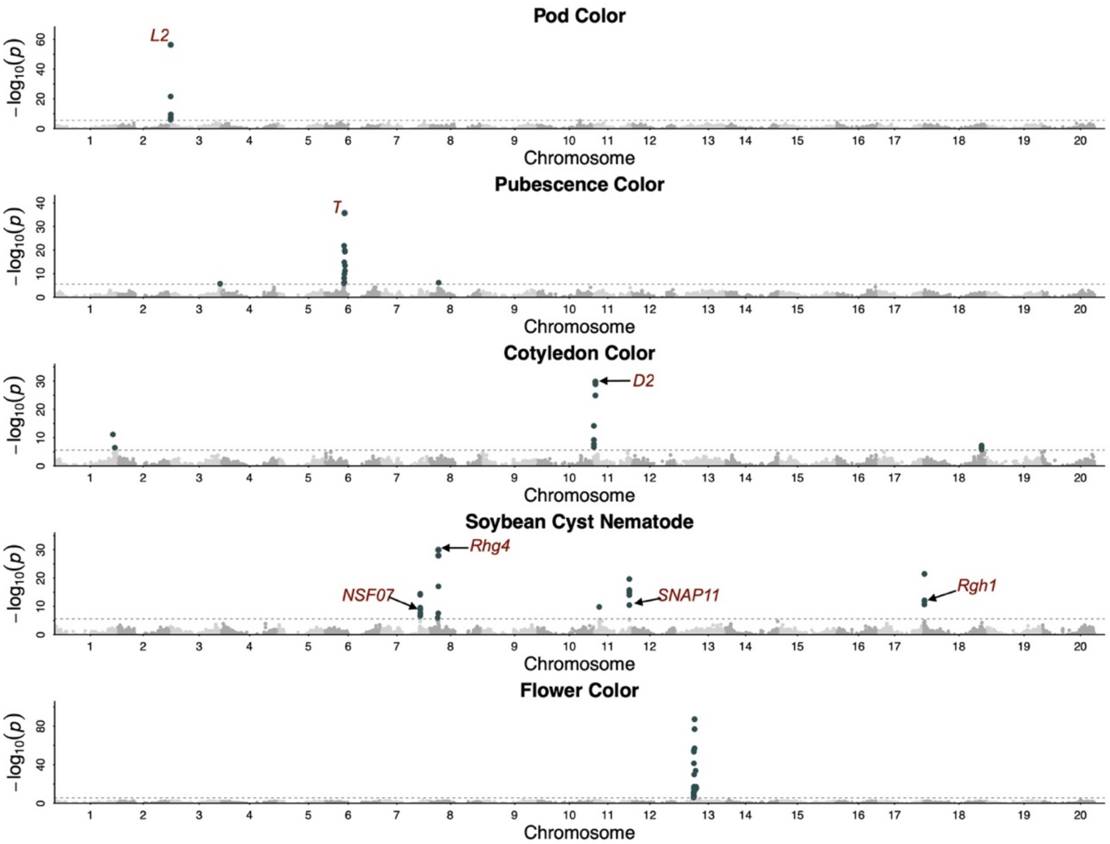


**Figure S6. TWAS FUSION for each of the five known traits.**

Manhattan plots of each FUSION TWAS for corresponding traits. Dashed line indicates the association cutoff. Each dot on the plot represents a feature, i.e., gene with *Cis*-eQTL or exon with *Cis*-exonQTL. The associated features that were found to be significant are highlighted in black, while non-significant ones are shown in grey. When known genes were identified as significantly associated, they were labeled with their most significant feature.

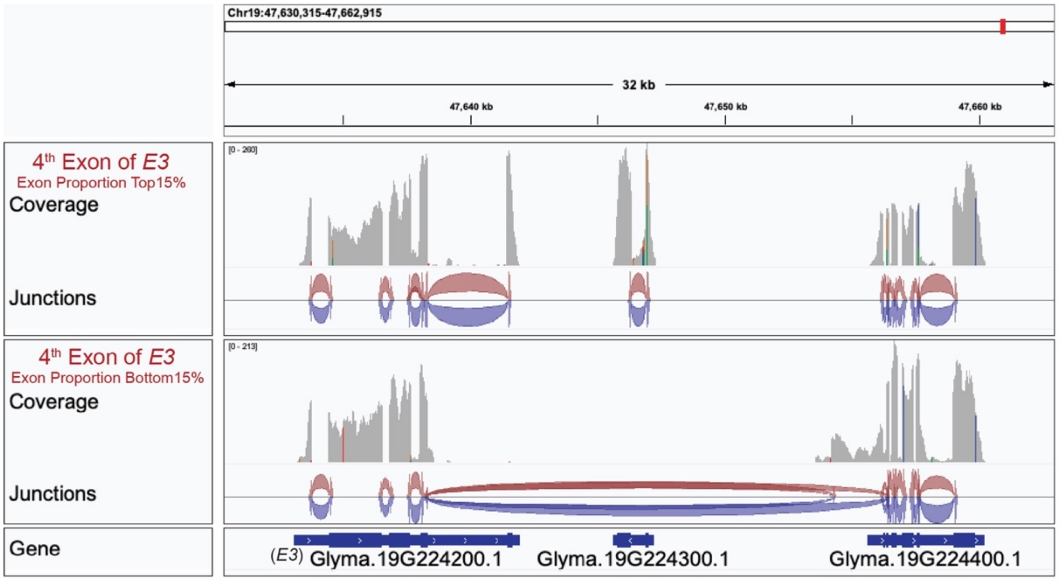


**Figure S7. Gene fusion of *E3* revealed with RNA-Seq data.**

Samples were categorized into the top 15% and bottom 15% on the reads proportion of 4th exon of *E3*. Five samples from each category were randomly selected and pooled for visualization. The reads coverage and junctions were shown with IGV for each pool.

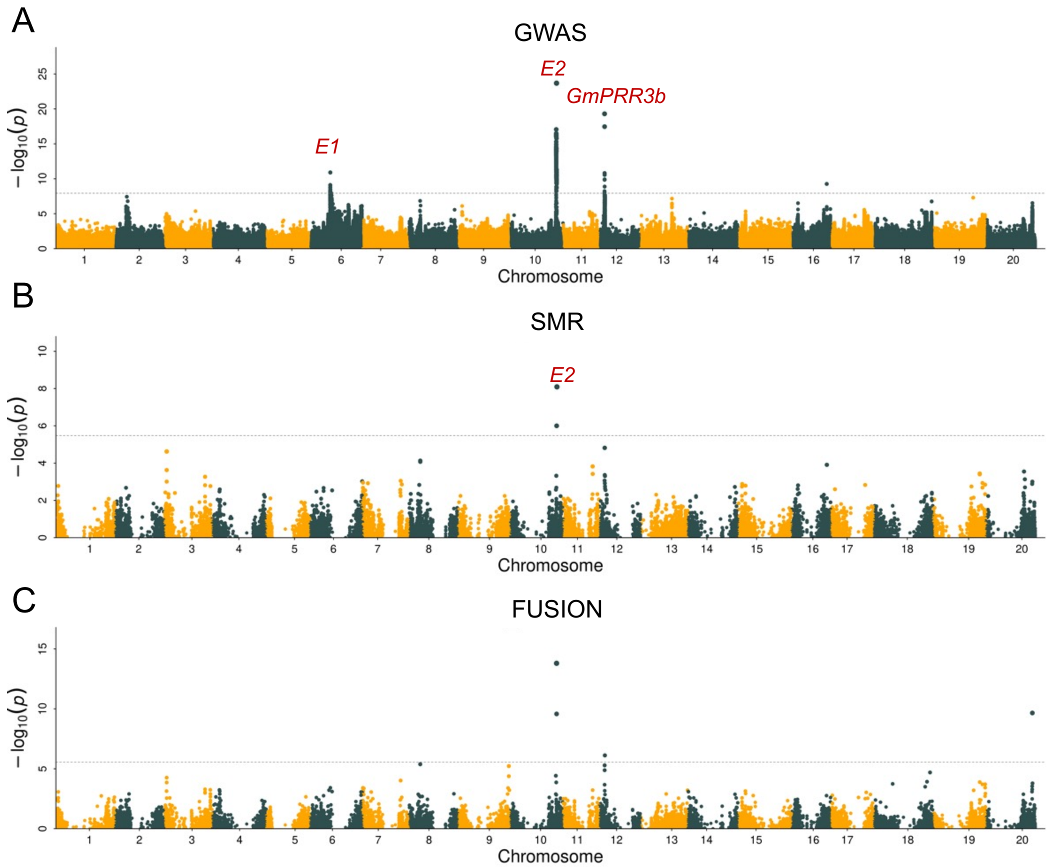


**Figure S8. Association analyses of flowering time.**

(**A**) A flowering time GWAS was conducted on 1,715 soybean cultivars with available flowering time and genotype data. Each dot represents one SNP. The three associated and already known loci were labeled in red. TWAS, integrating the GWAS results (**A**) with *Cis*-QTLs (eQTLs and exonQTLs) using different methods (SMR and FUSION), was shown in (**B-C**). (**B-C**), Each dot represents a feature, i.e., a gene with *Cis*-eQTL or an exon with *Cis*-exonQTL. The associated and known gene is labeled with its respective name.

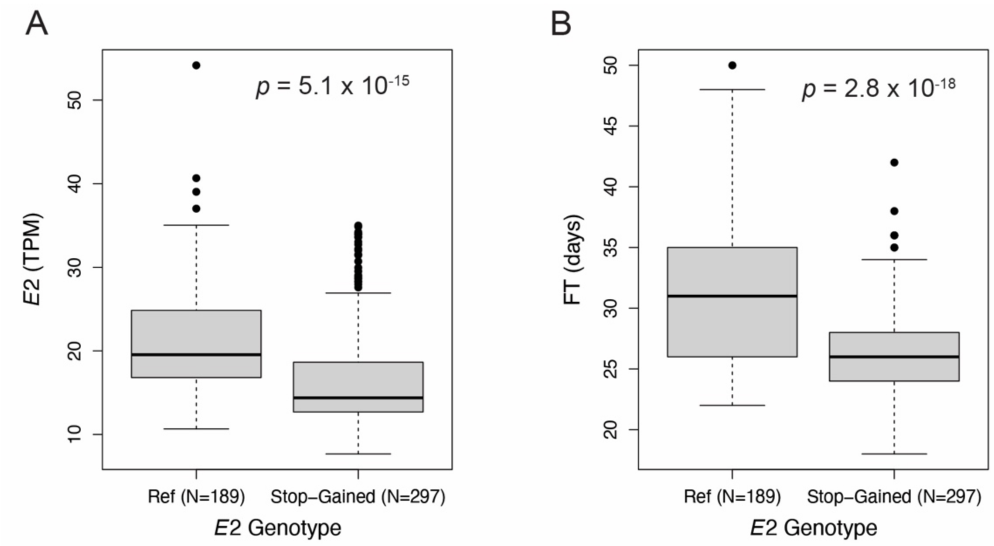


**Figure S9. Gene expression and flowering time for genotypes of the casual variant of *E2*.**

The presumed causal variant of *E2* was shared by *Cis*-eQTL of *E2* and flowering time GWAS as the leading SNP. The gene expression of *E2* (**A**) and flowering time (FT) (**B**) were shown for different genotype of the causal variant. The *p* values were resulted from Welch's two-sample t-test.

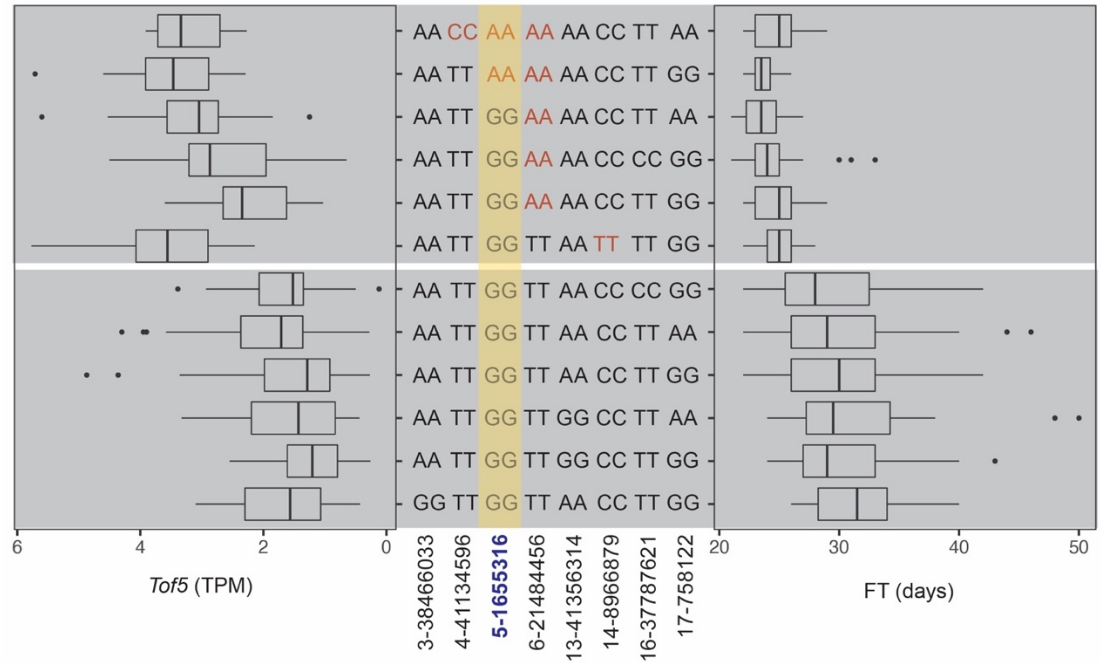


**Figure S10. Gene expression and flowering time for the different pseudo-haplotypes of eQTLs and exonQTL of *Tof5*.**

One *Cis*-eQTL (5-1655316), one *Trans*-exonQTL (3-38466033), and six *Trans*-eQTLs were identified for *Tof5*. Pseudo-haplotypes were created using leading SNPs of them, and pseudo-haplotypes with more than 10 individuals were retained. These pseudo-haplotypes were divided into two distinct categories based on either gene expression or flowering time. Four eQTLs, including the *Cis*-eQTL (highlighted in yellow), were shown with different genotypes in these two categories. The genotype associated with earlier flowering time was highlighted in red.

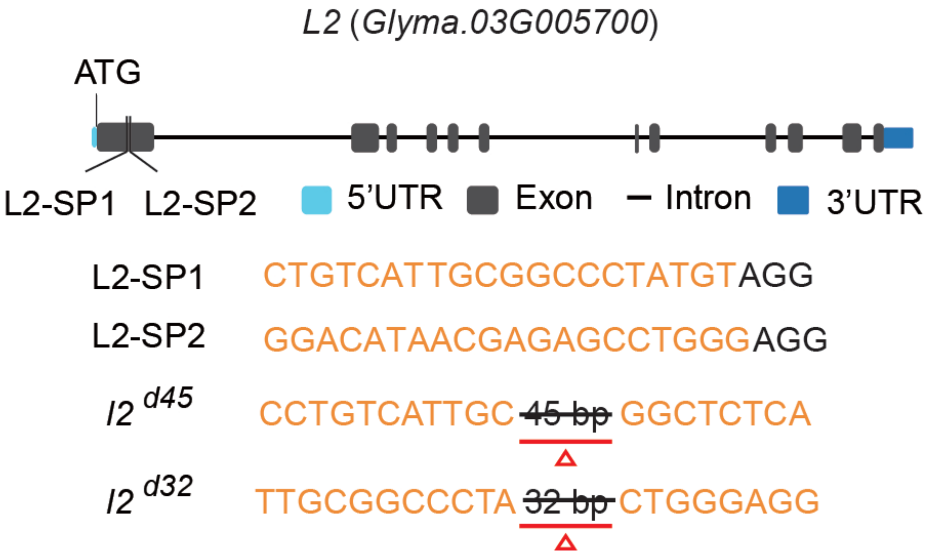


**Figure S11. The gRNAs for genome editing of *L2*.**

Two gRNAs, named L2-SP1 and L2-SP2, were designed for CRISPR/Case9 genome editing, and their targeting sites on the subject gene *L2* (*Glyma.03G005700*) were shown. The context sequencing of the two identified mutants were shown.
